# Prioritizing disease-related rare variants by integrating gene expression data

**DOI:** 10.1101/2024.03.19.585836

**Authors:** Hanmin Guo, Alexander Eckehart Urban, Wing Hung Wong

## Abstract

Rare variants, comprising a vast majority of human genetic variations, are likely to have more deleterious impact on human diseases compared to common variants. Here we present carrier statistic, a statistical framework to prioritize disease-related rare variants by integrating gene expression data. By quantifying the impact of rare variants on gene expression, carrier statistic can prioritize those rare variants that have large functional consequence in the diseased patients. Through simulation studies and analyzing real multi-omics dataset, we demonstrated that carrier statistic is applicable in studies with limited sample size (a few hundreds) and achieves substantially higher sensitivity than existing rare variants association methods. Application to Alzheimer’s disease reveals 16 rare variants within 15 genes with extreme carrier statistics. The carrier statistic method can be applied to various rare variant types and is adaptable to other omics data modalities, offering a powerful tool for investigating the molecular mechanisms underlying complex diseases.

## Introduction

Rare variants (minor allele frequency (MAF) < 1%) constitute a vast majority of human genetic variations^1,2^. They are on average more deleterious compared with common variants, and thus undergoes stronger selection and remains a low frequency in the general population. By analyzing large cohorts of whole genome sequencing/whole exome sequencing (WGS/WES) data, researchers have identified a handful of rare variants-trait association^3-6^ and shown that rare variants contribute to a large proportion of missing heritability that cannot be explained by common variants^7^.

Rare variants on average confer larger effects on gene expression and complex diseases and are easier to map to causal genes than common variants^8^. However, statistical power to identify disease-associated rare variants, especially for ultra-rare variants or even singletons, is limited, given that the sample size is not too high or the effect size is not too large. Variants collapsing methods (burden test, variance component test, omnibus test) are proposed to circumvent this obstacle^6,9-11^, which evaluate association for multiple variants in a biologically relevant region, such as a gene, instead of testing the effect of single variant. These methods work well under the assumption that multiple variants in a gene cumulatively contribute to the disease risk with each individual allele explaining only a small fraction of the cases, but resolution to pinpoint the risk variants may be diluted when the assumption is violated.

Complementary to genome sequencing assay, RNA-seq can quantitatively measure gene expression level and provide molecular cause of complex diseases, especially rare diseases. Previous studies have shown that rare variants are enriched near genes with aberrant gene expression^12-14^. We posit that those rare variants will be more prone to disease pathology. Specifically, for those rare variants with large functional impact, it will be likely to trigger some feedback in the regulatory network and lead to extreme expression change among disease-related genes. In this work, we propose carrier statistic, a statistical framework for prioritizing those rare variants with large functional consequence in diseased patients by integrating gene expression data. We demonstrate superior performance of our method through extensive simulations and application study to the Alzheimer’s disease, where given a limited sample size, existing rare variants association methods without functional gene expression data cannot provide positive findings.

## Results

### Method Overview

Our method stems from the expectation that diseased population shows enrichment in rare variants that have large impact on expression for disease-related genes. Suppose we have genotypes (e.g. variants call from WGS data) and gene expression measurements (e.g. reads count from RNA-seq data) for both diseased patients and healthy controls. For each rare variant-gene pair, we calculate the expression association z-score for rare variant carriers by using the gene expression from individuals without the variant as the null distribution (**Figure 1a**). The expression association z-score, which we term as carrier statistic, is calculated separately within the case and the control group. We only consider rare variant-gene pair wherein the variant is located within the exon of the gene throughout this study. The carrier statistic quantifies the degree to which the rare variant impacts gene expression level. We assume that most rare variants do not have a large impact on the gene function, so the distribution of carrier statistic will be centered around 0. Now, if a gene is relevant for the disease, then conditioning on having the disease will bias the sampling towards people carrying rare variants with large functional impact on the gene expression. Thus, the carrier statistics for disease-related rare variant-gene pairs will tend to be more extreme compared to those for non-related pairs in the case group (**Figure 1b**). We prioritize those rare variants and genes with outlier carrier statistic in the case group. False discovery rate (FDR) can be computed as the ratio of tail probability for carrier statistic between two groups (**Methods**).

**Figure 1.**
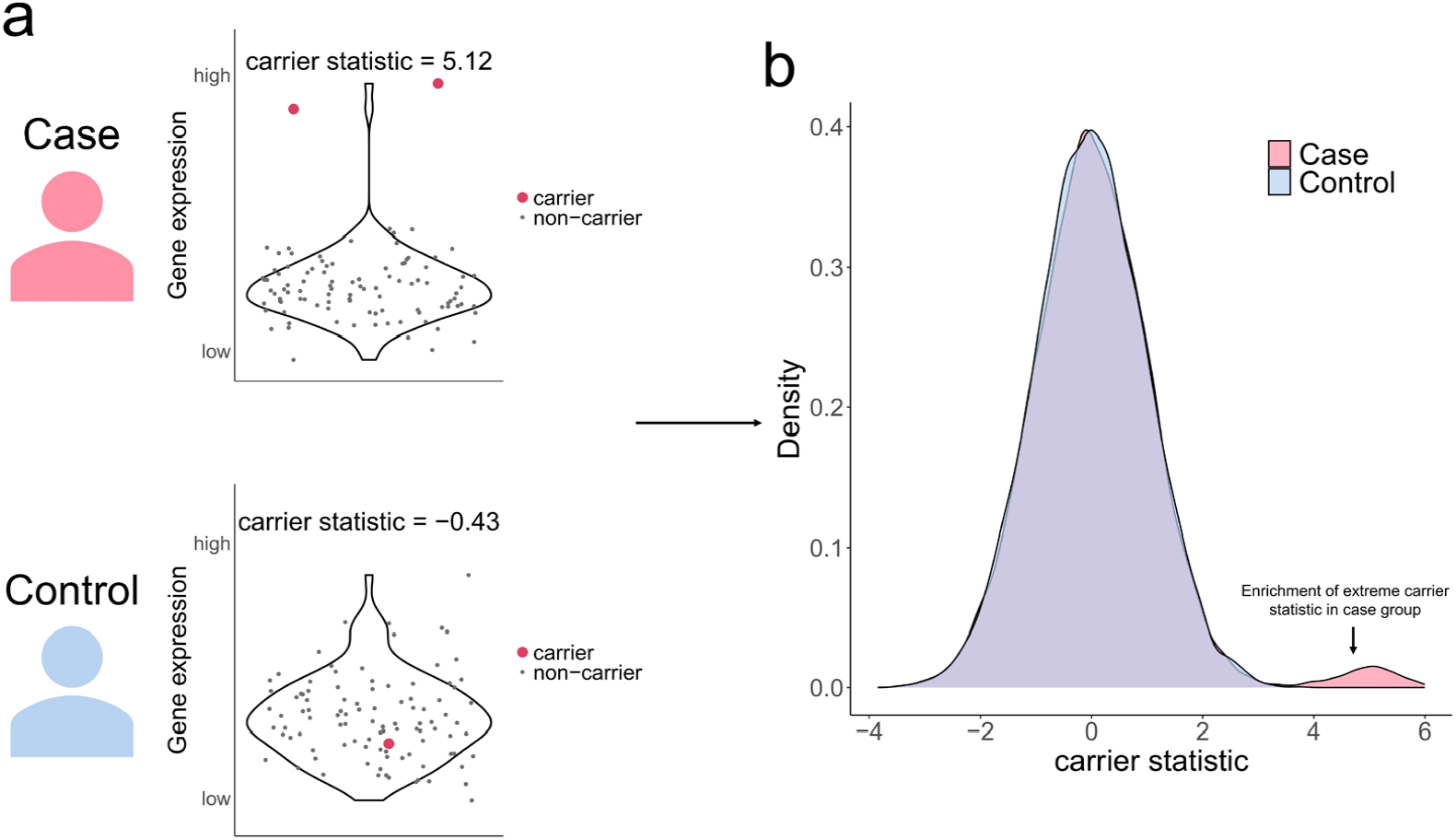
Carrier statistics for disease-related rare variant-gene pairs will tend to be more extreme compared to those for non-related ones in the case group.

### Simulation Results

We first carried out simulations to assess whether the carrier statistic-based method would produce false positive findings. We simulated genotypes based on whole exome sequencing (WES) data from Genome Aggregation Database (gnomAD)^2^ and simulated gene expression profiles based on RNA-seq data in whole blood tissue from the Genotype-Tissue Expression (GTEx) project (**Methods**). We perturbed the expression level of the causal genes for causal variants carriers by assuming that causal rare variants have large functional impact on disease-related genes. FDR for carrier statistic was well-calibrated in all simulation settings with varying penetrance of causal variant, prevalence in causal variant noncarriers, and number of causal variants per causal gene (**Supplementary Figure 1**). We also checked if using gene expression from noncarriers in two groups together rather than separately as null distribution will produce false positive findings, particularly when diseased patients have systematic change in gene expression profile from healthy controls (**Methods**). In this case, FDR showed substantial inflation for using gene expression from all noncarriers as null distribution, while using gene expression from noncarriers in the same group of carriers as null distribution consistently gave well-calibrated error rates (**Supplementary Figure 2**).

Next, we benchmarked the performance of carrier statistic with three existing rare variants association methods: burden test^9^, Sequence Kernel Association Test (SKAT)^10^, and SKAT-O^11^. Burden test counts the number of rare variants within a gene followed by the association test with the disease. SKAT computes a gene-level variance component score statistic which allows bidirectional effect of different variants. The unified test SKAT-O implements a linear combination of burden test statistic and variance component test statistic, which is preferred when the underlying genetic architecture of the disease is not known. While all four methods controlled FDR at the nominal level (**Supplementary Figure 1**), carrier statistic consistently achieved higher sensitivity than the three variants collapsing methods under various settings (**Figure 2**). It is worth noting that only a small proportion of case samples can be attributed to the causal variants in our simulated disease model, thus the enrichment of rare variants burden within causal genes in the case group over that in the control group was moderate and did not attain statistical significance (**Supplementary Tables 1-3**). However, carrier statistic could prioritize those causal rare variants with large functional impact by integrating information from gene expression data.

**Figure 2.**
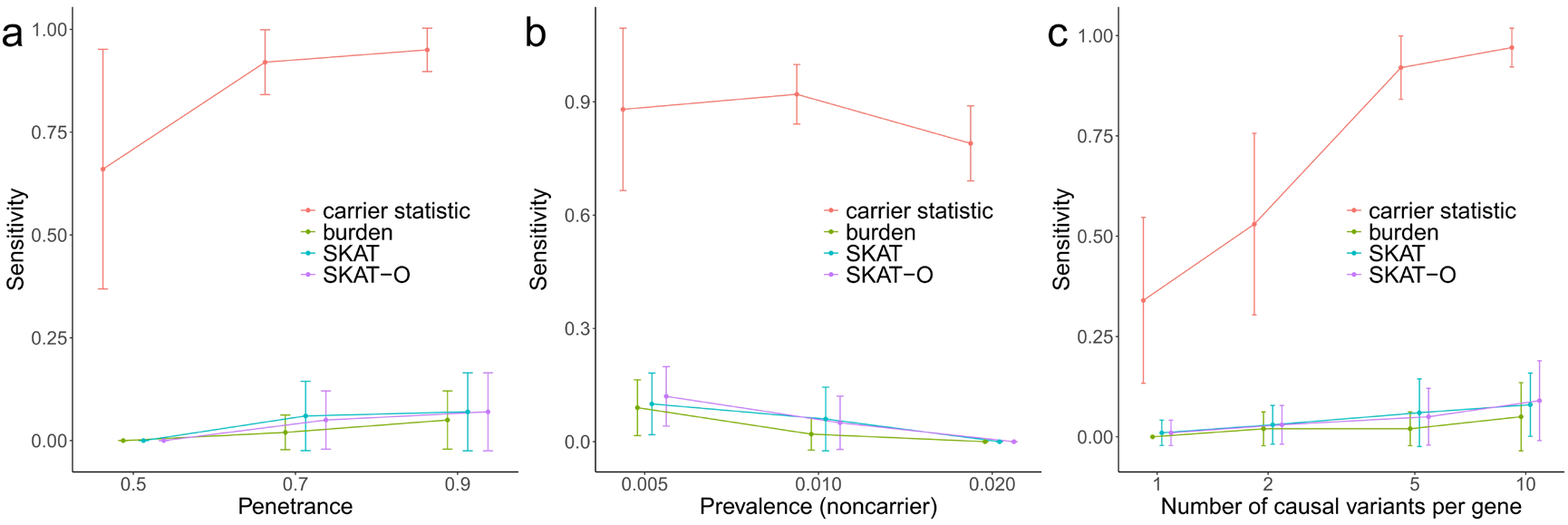
Carrier statistic achieves higher sensitivity than variants collapsing methods in simulations with varying (a) penetrance of causal variant, (b) prevalence in causal variant noncarriers, and (c) number of causal variants per causal gene.

### Application to Alzheimer’s disease

Alzheimer’s disease is highly heritable, with estimated heritability up to ∼60%-80% based on twin studies^15^. Large scale GWASs have identified multiple loci contributing to Alzheimer’s disease, but the genetic variance explained by the significant loci is far from the level suggested by the disease heritability^16^. Additionally, there is limited understanding regarding the molecular mechanism through which GWAS significant variants affect the disease, with the exception of the well-known APOE locus. To investigate whether rare variants (single nucleotide variants [SNVs] and short indels) confer functional consequences in Alzheimer’s disease, we applied carrier statistic to a harmonized multi-omics dataset (*n*_*case*_ = 444, *n*_*control*_ = 234) consisting of WGS and RNA-seq from four aging cohort studies: the Religious Orders Study (ROS) and Memory and Aging Project (MAP), the Mount Sinai Brain Bank (MSBB), and the Mayo Clinic (**Methods**). We found significant excess of large carrier statistic in the diseased patients (**Figure 3**). Controlling FDR with a cutoff of 0.2, we prioritized 16 rare variants within 15 genes with large carrier statistic in the case group (**Table 1**), implicating those variants may contribute to Alzheimer’s disease through up-regulating gene expression in the brain.

**Figure 3.**
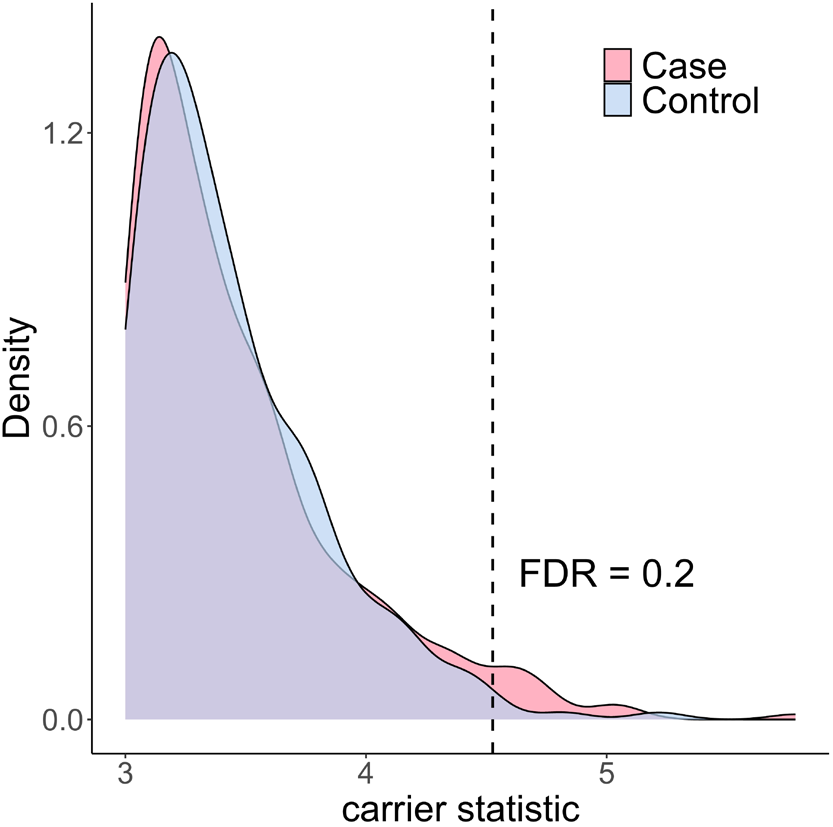
Alzheimer’s disease patients show significant excess of large carrier statistic.

**Table 1.** 16 rare variants within 15 genes with large carrier statistic in the Alzheimer’s disease patients.

| CHR | POS | REF | ALT | SNP | Gene | Carrier statistic |
| --- | --- | --- | --- | --- | --- | --- |
| 14 | 31344166 | G | A | rs200935305 | COCH | 5.78 |
| 10 | 71716481 | G | A | rs537156091 | COL13A1 | 5.09 |
| 3 | 147108829 | T | TGTAGCCC | rs765482543 | ZIC4 | 5.02 |
| 15 | 63845640 | C | T | rs971269728 | USP3 | 5.00 |
| 15 | 81565527 | G | T | rs188062582 | IL16 | 4.77 |
| 10 | 115991333 | A | G | rs867202477 | TDRD1 | 4.74 |
| 6 | 37614984 | C | T | rs755089010 | MDGA1 | 4.71 |
| 1 | 29445783 | A | AT | rs1210817818 | EPB41 | 4.69 |
| 5 | 161113972 | A | G | rs966622465 | GABRA6 | 4.69 |
| 15 | 32907368 | GCCTTCAGC | G | rs1377672532 | ARHGAP11A | 4.62 |
| 7 | 31747403 | G | A | rs766182885 | PPP1R17 | 4.61 |
| 18 | 22804990 | G | A | rs770368967 | ZNF521 | 4.61 |
| 6 | 34028294 | C | T | rs556027021 | GRM4 | 4.58 |
| 1 | 231356659 | G | A | rs528099262 | TRIM67 | 4.56 |
| 6 | 37665148 | A | G | rs545009624 | MDGA1 | 4.55 |
| 6 | 123046921 | C | T | rs1016716974 | PKIB | 4.53 |

To benchmark the performance of carrier statistic with existing methods, we also applied burden test, SKAT, SKAT-O to the same Alzheimer’s disease dataset. These three variants collapsing methods did not identify any significant genes (FDR < 0.2), possibly due to insufficient sample size. To further evaluate the performance of carrier statistic, we assessed the enrichment of rare variants burden within the top prioritized genes in case group compared to that in controls. Among the top 100 genes with largest carrier statistic, 67 genes have fold enrichment larger than 1, 32 genes have fold enrichment larger than 4/3, while only 6 genes have fold enrichment smaller than 3/4 (**Supplementary Figure 3**). As demonstrated in the simulations, enrichment of rare variants burden within those genes was moderate and did not pass significance threshold by the variants collapsing methods.

The carrier statistic method prioritized a handful of significant genes that may shed light on the genetic etiology of Alzheimer’s disease (**Figure 4**). *COCH* has the largest carrier statistic of 5.78. Missense mutations within this gene were found to cause the late-onset DFNA9 deafness disorder^17,18^. Furthermore, deposits of Cochlin encoded by *COCH* is associated with age-related glaucomatous trabecular meshwork but absent in healthy controls. Additionally, SNPs inside *COCH* are associated with cortical thickness^19^, changes in which through neuroimaging techniques is commonly used in early detection and monitoring of Alzheimer’s disease progression^20,21^. Gene *ARHGAP11A* has a carrier statistic of 4.62. Transcribed mRNAs of the gene subcellularly localize and are locally translated in radial glia cells of human cerebral cortex and further regulate cortical development^22^. More importantly, *ARHGAP11A* may contribute to Alzheimer’s disease pathology by mediating Amyloid-β generation and Amyloid-β oligomer neurotoxicity^23^. *PPP1R17* (carrier statistic = 4.61) functions as a suppressor of phosphatase complexes 1 (PP1) and 2A (PP2A). A recent mice study suggests that hypothalamus neurons with *PPP1R17* expression are associated with synaptic dysfunction and WAT impairment, which can further delay aging^24^. Interestingly, *PPP1R17* is involved in human-specific cortical neurodevelopment regulated by enhancers in human accelerated regions^25^. *ZIC4* (carrier statistic = 5.02), also known as zinc finger protein of the cerebellum 4, plays an important role in the embryonal development of the cerebellum. Heterozygous deletions encompassing the *ZIC4* locus are associated with a rare congenital cerebellar malformation known as the Dandy–Walker malformation^26^. Notably, mutations in proximity to the *ZIC4* loci are implicated in multiple system atrophy, a rare neurodegenerative disease^27^. Large scale GWAS study of brain morphology has also identified associations with *ZIC4*, underscoring its significance in diverse neurological processes^28^. *MDGA1* (carrier statistic = 4.71) encodes a glycosylphosphatidylinositol (GPI)-anchored cell surface glycoprotein. It has been reported that *MDGA1* can contribute to cognitive deficits through altering inhibitory synapse development and transmission in the hippocampus^29^. Of note, *MDGA 1* is one of the 96 genes from the Olink neurology panel with established links to neurobiological processes and neurological diseases.

**Figure 4.**
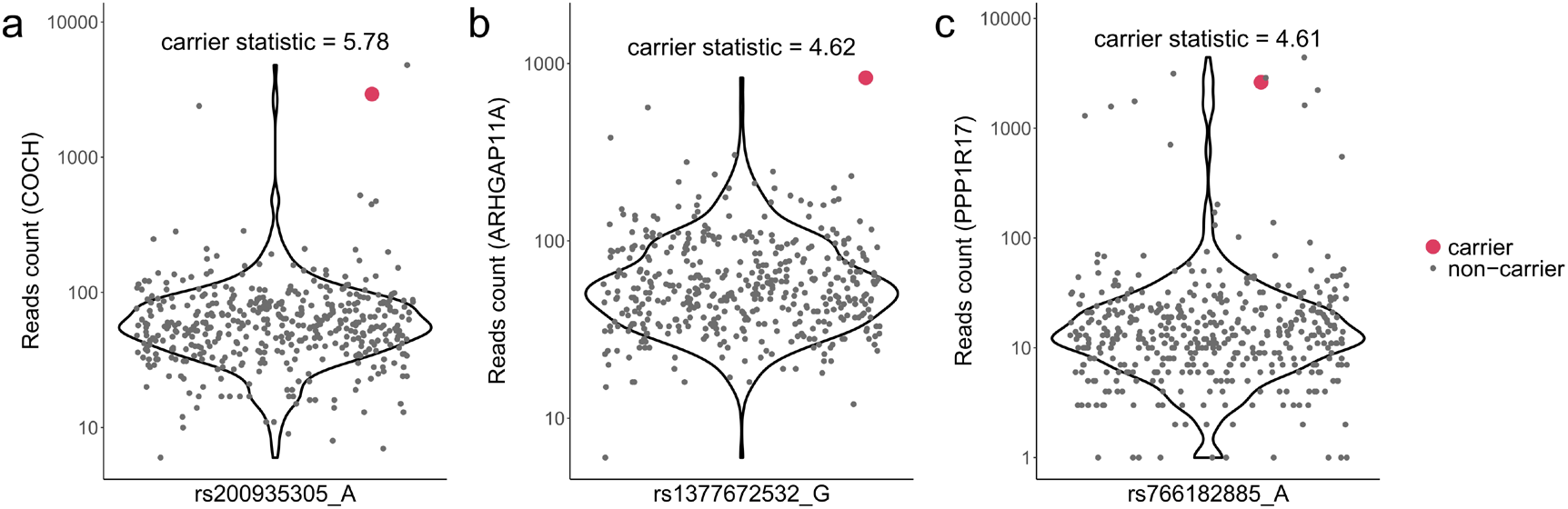
Rare variant carriers show outlier expression level for (a) *COCH*, (b) *ARHGAP11A*, and (c) *PPP1R17*. Pseudo count 1 was added to the RNA reads count for visualization purpose. Y-axis is on log scale.

## Discussion

We presented carrier statistic, a statistical framework to perform multi-omics data analysis, for prioritization of disease-related rare variants and their regulated genes. Through simulations and analyses of real multi-omics dataset, we demonstrated that carrier statistic overcomes sample size limitation and achieves substantial gain in statistical power compared to existing variants collapsing methods. The superior performance of carrier statistic can be attributed to incorporation of functional gene expression data, which allows quantitatively measuring the impact of rare variants that cannot be determined by looking at the variants alone. We applied carrier statistic to Alzheimer’s disease and highlighted several novel risk genes, providing insights into the molecular etiology of the complex disease.

Carrier statistic serves as a general approach, for the first time, to study how rare variants affect complex disease through mediating gene expression. There exist several methods such as transcriptome-wide association study (TWAS)^30,31^ or colocalisation^32,33^ that can also perform integrative analysis across multiple data modalities (genotype, gene expression, and phenotype). However, those methods focus exclusively on effects of common SNPs and will have limited power for rare variants. In addition to SNVs and short indels that we included in this study, the statistical framework can be also applied to other types of rare variants (simple structural variants [SVs], complex SVs, mobile element insertions, tandem repeat expansions), which in general have larger effect size than SNVs^34^. Finally, carrier statistic can be adapted to other omics data, such as epigenomics and proteomics. As multi-omics data accumulates complementary to genome sequencing assay, we anticipate that carrier statistic will be an effective approach to dissect the molecular mechanism of complex diseases.

## Methods

### Carrier statistic

For each rare variant-gene pair (the variant is located within the exon of the gene), we used the expression of that gene in the rare variant noncarriers as the null distribution and computed a z-score for each rare variant carrier, then average over carriers of that variant. The carrier statistic was computed separately within the case group and the control group. Rare variants were defined as SNVs and short indels whose allele count was no larger than 5 within the case group or within the control group. Therefore, the rare variants and thus the number of carrier statistics are not the same between two groups. The carrier statistic quantifies the degree to which the rare variant impacts gene expression level. We assume that diseased population shows enrichment in rare variants that have large impact on expression for disease-related genes, thus there will also be enrichment of extreme carrier statistic for disease-related rare variant-gene pairs in the case group. We prioritize those rare variants and genes with outlier carrier statistic in the case group. For rare variant-gene pairs with positive carrier statistic, false discovery rate at a given threshold of carrier statistic, denoted by *z*_0_, can be computed as 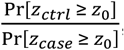, where *z*_*case*_ and *z*_*crtl*_ denote the carrier statistic in the case group and control group, respectively. Duplicative carrier statistics were removed (i.e. multiple rare variants occurring in the same individuals are counted as the same rare variant). Similarly, for rare variant-gene pairs with negative carrier statistic, false discovery rate at threshold of *z*_0_ can be computed as 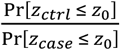.

### Simulations

We simulated genotypes for a large population consisting of 125,748 individuals based on the alternative allele count from 125,748 exomes in the gnomAD v2.1.1 dataset^2^. Only exonic variants that passed all variant filters in the gnomAD dataset were retained. Then we simulated gene expression data for the large population as follows. We first simulated background gene expression profile for these 125,748 individuals while matching the mean and standard deviation of normalized gene expression in the reference expression dataset. In this study, we used *log*_2_(reads count + 1) as normalized gene expression and RNA-seq in the whole blood tissue from GTEx project v8 as the reference expression dataset^35^. Genes whose median number of reads count in the large population < 10 were removed. We randomly selected *m* causal genes and *l* causal variants for each causal gene, where *m* was set as 10 and *l* was set to vary from 1 to 10. For each causal gene, we perturbed the gene expression for causal variant carriers by *z * sd* fold, where *z* was set to vary from 1 to 5 and *sd* was the standard deviation of normalized gene expression in the reference dataset. Next, we simulated the disease status for the large population by assuming penetrance of causal variant as *p*_*carrier*_ and prevalence in causal variant noncarrier as *p*_*noncarrier*_. Here *p*_*carrier*_ varied from 0.5 to 0.9 and *p*_*noncarrier*_ varied from 0.005 to 0.02. Finally, we randomly sampled 500 cases and 500 controls from the affected and nonaffected population respectively to mimic the sample recruitment procedure in the disease study. Each simulation setting was repeated for 100 times.

We evaluated the performance of different methods using two metrics: FDR and sensitivity. FDR was defined as the proportion of falsely identified genes among all identified ones. If no gene was identified then FDR was set as 0. Sensitivity was defined as the proportion of truly identified genes among all underlying causal genes.

Note that carrier statistic was computed by using gene expression from rare variant noncarriers in the same group (i.e. case or control) as the carriers as null distribution. We also checked if using gene expression from noncarriers in both case group and control group as the null distribution will produce false positive findings. We perturbed expression level for all genes in the case group. In this case, FDR showed substantial inflation for using gene expression from all individuals in two groups as null distribution, especially when there is large systematic difference in the transcriptome between two groups (**Supplementary Figure 1b**). On the contrary, using gene expression from rare variant noncarriers in the same group of carriers as null distribution consistently controlled FDR at the nominal level (**Supplementary Figure 1a**).

### Implementation of different methods

Variants collapsing methods were performed using the R package SKAT v.2.2.5. All the parameters were set as default value. Both common and rare variants were included in the analysis.

### Multi-omics data analysis for Alzheimer’s disease

WGS data for the four aging cohorts (ROS/MAP, MSBB, and the Mayo Clinic) were obtained from the Whole Genome Sequence Harmonization Study (Synapse ID: syn22264775). RNA-seq data for the same four cohorts were downloaded from the RNAseq Harmonization Study (syn21241740). Only white people with both WGS and RNA-seq data were included in the analysis.

We determined disease status following description in previous publications^36,37^. For the ROS/MAP cohorts, individuals with a Braak neurofibrillary tangle score *≥* 4, a CERAD neuritic and cortical plaque score ≤ 2, and a cognitive diagnosis of probable Alzheimer’s disease with no other causes (cogdx = 4) were classified as cases, while individuals with a Braak score ≤ 3, a CERAD score *≥* 3, and a cognitive diagnosis of no cognitive impairment (cogdx = 1) were classified as controls. For MSBB, individuals with a Braak score *≥* 4, a CERAD score *≥* 2, and a Clinical Dementia Rating (CDR) score *≥* 1 were classified as cases, while individuals with a Braak score ≤ 3, a CERAD score ≤ 1, and a CDR score ≤ 0.5 were classified as controls. For the Mayo Clinic cohort, individuals with a Braak score *≥* 4 and a CERAD score *≥* 2 were classified as cases, while individuals with a Braak score ≤ 3 and a CERAD score ≤ 1 were classified as controls. Of note, definition of CERAD score in the ROS/MAP cohort is different from that in the MSBB and the Mayo Clinic cohorts. After harmonization across cohorts, 444 cases and 234 controls in total were identified and used for downstream analysis.

Next, we performed quality control on the WGS and RNA-seq data. For WGS data, only exonic variants with missing genotypes < 10% were retained. For RNA-seq data, we selected prefrontal cortex as the target brain tissue. If a donor does not have RNA-seq in the prefrontal cortex, then RNA-seq in other tissues will be used based on the following order: dorsolateral prefrontal cortex > posterior cingulate cortex > head of caudate nucleus in the ROS/MAP cohort, prefrontal cortex > frontal pole > superior temporal gyrus > inferior frontal gyrus > parahippocampal gyrus in the MSBB cohort, and temporal cortex > cerebellum in the Mayo Clinic cohort. Genes with zero reads count in more than 10% of samples or with median reads count < 10 across samples were excluded. *log*_2_(reads count + 1) was used as normalized gene expression. Then we applied carrier statistic to perform downstream analysis.

## Data availability

The WGS data in the Whole Genome Sequence Harmonization Study (https://www.synapse.org/#!Synapse:syn22264775) and the RNA-seq data in the RNAseq Harmonization Study (https://www.synapse.org/#!Synapse:syn21241740) are publicly available on the AD Knowledge Portal platform through completion of a data use certificate. The gnomAD v2.1.1 data consisting of 125,748 exomes were downloaded from https://gnomad.broadinstitute.org/. The gene expression data (gene reads count) from GTEx project version 8 were downloaded from the GTEx Portal, https://gtexportal.org/home/downloads/adult-gtex/bulk_tissue_expression.

## Acknowledgements

We thank Dr. Bo Zhou and Dr. Hua Tang for discussion and providing feedbacks.

This work was partially supported by NIH grants R01 HG010359, R01 MH116529, and P50 HG007735.

## Author contributions

W.H.W. and A.E.U. supervised the study. H.G. and W.H.W. designed the idea. H.G. performed data analysis. H.G., A.E.U., and W.H.W. interpreted the results. H.G. and W.H.W. wrote the manuscript.

