## Supplementary Figures for "Prioritizing disease-related rare variants by integrating gene expression data"

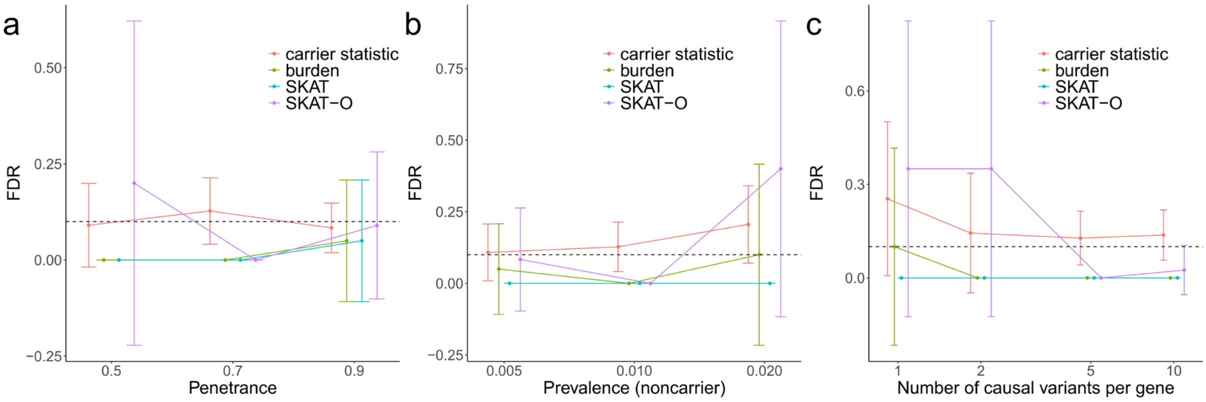


**Supplementary Figure 1. FDR for carrier statistic, burden test, SKAT, and SKAT-O in simulations with varying (a). penetrance of causal variant, (b) prevalence in causal variant noncarriers, and (c) number of causal variants per causal gene.** Error bar shows standard deviation across 100 simulation repeats.

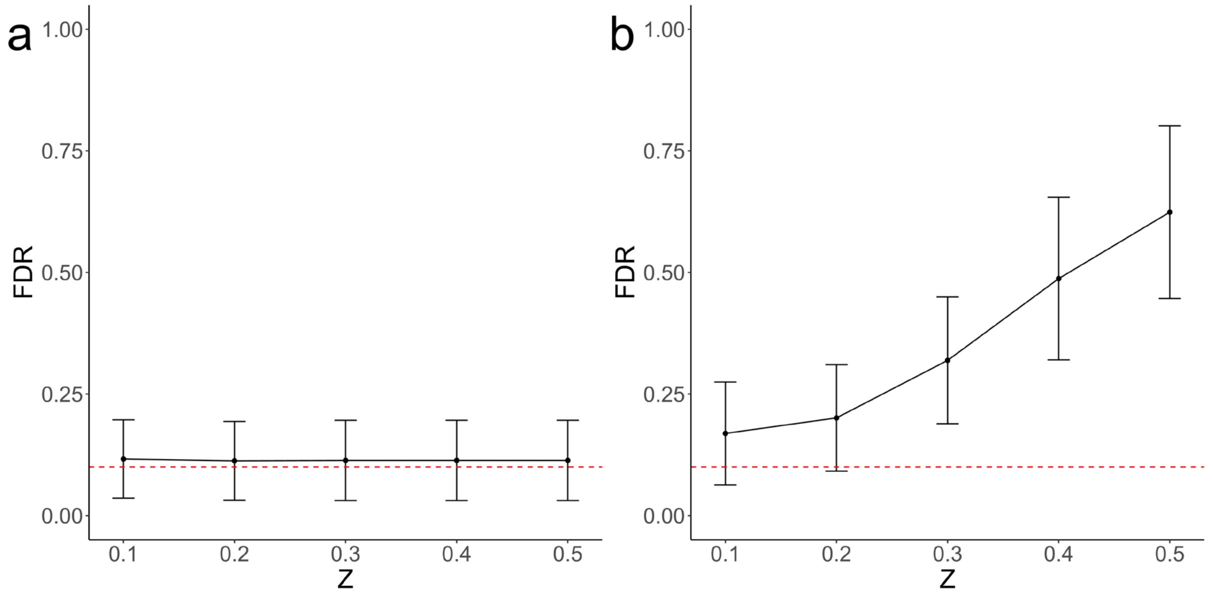


**Supplementary Figure 2. (a). FDR is well-calibrated for using gene expression from noncarriers in the same group of carriers as null distribution. (b). FDR showed substantial inflation for using gene expression from all noncarriers as null distribution.** Z quantifies the level of systematic difference in the transcriptome between case group and control group. Error bar shows standard deviation across 100 simulation repeats.

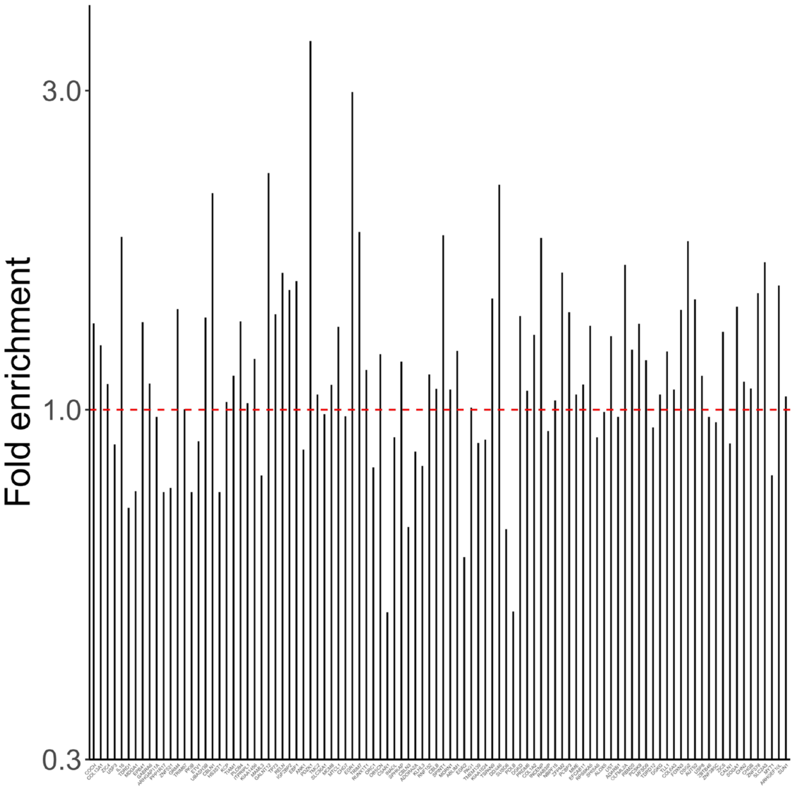


**Supplementary Figure 3. Enrichment of rare variants within top 100 genes with largest carrier statistic in Alzheimer's disease patients.** Genes were ranked according to decreasing order of carrier statistic. Fold enrichment was defined as the ratio of rare variants burden within the gene in case group compared to that in the control group.
