## Supplementary Tables for "Prioritizing disease-related rare variants by integrating gene expression data"

STable 1. Rare variants burden in simulated disesae model with varying penetrance of causal variant

| Penetrance of causal variant | Number of case samples carrying causal variant | Number of control samples carrying causal variant | Number of rare variants in causal genes per case sample | Number of rare variants in causal genes per control sample |
|---|---|---|---|---|
| 0.5 | 35.1 | 0.3 | 0.22 | 0.16 |
| 0.7 | 60 | 0.1 | 0.27 | 0.16 |
| 0.9 | 69.2 | 0 | 0.29 | 0.15 |

STable 2. Rare variants burden in simulated disesae model with varying prevalence in causal variant noncarriers

| Prevalence (causal variant noncarriers) | Number of case samples carrying causal variant | Number of control samples carrying causal variant | Number of rare variants in causal genes per case sample | Number of rare variants in causal genes per control sample |
|---|---|---|---|---|
| 0.005 | 97.7 | 0.4 | 0.37 | 0.17 |
| 0.01 | 60 | 0.1 | 0.27 | 0.16 |
| 0.02 | 23.8 | 0.1 | 0.17 | 0.14 |

STable 3. Rare variants burden in simulated disesae model with varying number of causal variants per causal gene

| Number of causal variants per causal gene | Number of case samples carrying causal variant | Number of control samples carrying causal variant | Number of rare variants in causal genes per case sample | Number of rare variants in causal genes per control sample |
|---|---|---|---|---|
| 1 | 8 | 0 | 0.15 | 0.13 |
| 2 | 25 | 0.1 | 0.17 | 0.12 |
| 5 | 60 | 0.1 | 0.27 | 0.16 |
| 10 | 102.8 | 0.5 | 0.37 | 0.18 |
